# A DNA demethylase overexpression promotes early apical bud maturation in poplar through the biosynthesis and accumulation of flavonoids

**DOI:** 10.1101/114652

**Authors:** Daniel Conde, Alicia Moreno-Cortés, Christopher Dervinis, José M. Ramos-Sánchez, Matias Kirst, Mariano Perales, Pablo González-Melendi, Isabel Allona

## Abstract

The transition from active growth to dormancy is critical for the survival of perennial plants. We identified a *DEMETER-like* (*CsDML*) cDNA from a winter-enriched cDNA subtractive library in chestnut *(Castanea sativa* Mill.), an economically and ecologically important species. Next, we characterized this DNA demethylase and its putative orthologue in the more experimentally tractable hybrid poplar *(Populus tremula × alba),* under the signals that trigger bud dormancy in trees. We performed phylogenetic and protein sequence analysis, gene expression profiling and 5mC immunodetection studies to evaluate the role of CsDML and its homologue in poplar, PtaDML6. Transgenic hybrid poplars overexpressing *CsDML* were produced and analyzed. Short days (SD) and cold temperatures induced *CsDML* and *PtaDML6.* Overexpression of *CsDML* accelerated SD-induced bud formation, specifically from stage 1 to 0. Bud acquired a red-brown coloration earlier than wild type (WT) plants, alongside with the upregulation of flavonoid biosynthesis enzymes and accumulation of flavonoids in the SAM and bud scales. Our data shows that the *CsDML* gene induces bud formation needed for the survival of the apical meristem under the harsh conditions of winter. This study provides *in planta* evidence implicating chromatin remodeling by DNA demethylation during SD induction of bud maturation through the induction of flavonoids biosynthesis.

## INTRODUCTION

Coordinating growth and reproduction with the environment is essential for the survival of perennial plants in temperate and boreal latitudes (Peñuelas and Filella, 2001). In these regions, deciduous and periodic growth habits evolved into a single trait known as winter dormancy. Winter dormancy begins with growth cessation through the arrest of meristems activity and subsequent enclose of the apical meristem into a dormant winter bud, a process called bud set (Rohde and Bhalerao, 2007; Paul et al., 2014). This process is required for the establishment of dormancy in the meristem, although considerable time may elapse before full winter dormancy is achieved (Rohde et al., 2002; Rohde and Bhalerao, 2007). For some perennial plants such as poplar, the perception of SD is sufficient to promote cessation of growth and bud formation (Ding and Nilsson, 2016). These processes may be positively influenced by synergic crosstalk between SD and temperature decrease in autumn (Rohde et al., 2011a).

Early insights into the molecular mechanisms of growth cessation and bud set induced by SD were gained through the analysis of the poplar homologues of *Arabidopsis (Arabidopsis thaliana* L.) *CONSTANS (CO)* and *FLOWERING LOCUS T (FT)* (Böhlenius et al., 2006). Downregulation of the *FT2* gene by SD was shown to be an early event promoting growth cessation (Böhlenius et al., 2006; Hsu et al., 2011). Downstream of *FT2,* and during the subsequent downregulation of *FLOWERING LOCUS D-LIKE1 (FDL1),* MADS box transcription factors Like-APETALA1 (LAP1) and AINTEGUMENTA LIKE1 (AIL1) were also found to play a role in growth cessation under SD conditions in poplar apices (Karlberg et al., 2010; Petterle et al., 2013; Azeez et al., 2014; Tylewicz et al., 2015). Phytochrome photoreceptors and circadian clock core components upstream of these genes have also been implicated in the early regulation of growth cessation by SD (Ibáñez et al., 2010; Kozarewa et al., 2010; Ding and Nilsson, 2016). While shoot elongation ceases, apical buds develop. During this process, leaf primordia become bud scales enclosing the apical meristems and preformed embryonic shoots. These bud scales accumulate phenolic compounds, leading to their characteristic red-brown color (Rohde et al., 2002). ABI3 is involved in this processes and ethylene resistant birch shows an altered bud development (Rohde et al., 2002; Ruonala et al., 2006). SD signals also influence growth by modulating gibberellic acid and indole-acetic acid pathways (Eriksson et al., 2000; Petterle et al., 2013; Zawaski and Busov, 2014).

Several transcriptome studies in *Populus spp.* and other perennial plants have correlated the expression patterns of genes linked to DNA methylation with the transition from active growth to dormancy of the apical and stem meristems (Karlberg et al., 2010; Rios et al., 2014; Shim et al., 2014; Howe et al., 2015). Genomic DNA methylation is known to be involved in numerous aspects of chromatin function, particularly gene expression (Law and Jacobsen, 2010). In plants, three classes of DNA methyltransferases catalyze 5-methyl-cytosine methylation (5mC) in symmetrical (CG or CHG) and nonsymmetrical (CHH) contexts (H represents A, T or C) (Zhang et al., 2010). However, the opposite process, regulation of 5mC DNA demethylation, is less well understood. DNA methylation may take place passively through the synthesis of unmethylated strands during DNA replication (Law and Jacobsen, 2010), or actively through the actions of enzymes that remove methylated bases. Plants have evolved specific enzymes to remove methylated cytosines, and these include a subgroup of DNA glycosylase-lyases known as DEMETER LIKE DNA demethylases (DMLs). Methylome studies performed in *Arabidopsis* and *Populus trichocarpa* (Torr & Gray) have demonstrated that DNA methylation in promoter and upstream sites near the transcriptional start site (TSS) suppress gene expression (Zhang et al., 2006; Vining et al., 2012; Lafon-Placette et al., 2013; Liang et al., 2014). In *Arabidopsis,* active DNA demethylation by DMLs reduces extensive DNA methylation near TSS and the 3' end of the transcribed region in hundreds of discrete *loci* of the genome (Penterman et al., 2007; Zhu et al., 2007; Ortega-Galisteo et al., 2008). Prior work in *Arabidopsis,* tomato *(Solanum lycopsersicum* L.) and tobacco *(Nicotiana tabacum* L.) has assigned DMLs a main role as transcriptional activators of their target genes. Indeed, gene activation by DMLs is necessary for the response of plants to biotic and abiotic stresses (Yu et al., 2013; Le et al., 2014; Bharti et al., 2015) and for plant development (Gehring et al., 2006; Liu et al., 2015). Epigenetic regulation of adaptive responses of forest tree species to the environment, such as changes in DNA methylation, have been implicated in the switch from active growth to dormancy (Santamaría et al., 2009; Conde et al., 2013; Zhu et al., 2013; Kumar et al., 2016), possibly one of the most critical developmental transitions for the survival of perennial plants. However, functional studies have not confirmed this hypothesis. Here we report the identification, phylogenetic and expression analyses and phenological studies of a chestnut 5mC DNA demethylase that triggers apical bud maturation under SD conditions in poplar.

## RESULTS

### Identification of a *DEMETER-LIKE* gene during autumn in *Castanea sativa*

To identify candidate genes regulating the winter dormancy process, we selected a cDNA showing similarity to a DNA glycosylase that was preferentially expressed at the beginning of autumn (Fig. S2A). InterPro analysis of the full-length sequence revealed the presence of a conserved HhH-GPD domain (helix-hairpin-helix and Gly/Pro rich loop followed by a conserved aspartate). The HhH-GPD domain belongs to the superfamily of the DNA glycosylases, which excise oxidized and methylated bases, correcting base pairing mismatches (Denver et al., 2003). Phylogenetic analyses of the *Arabidopsis* and poplar homologues of this chestnut DNA glycosylase showed that the protein clusters in the clade of 5mC DNA glycosylases (5mC), closer to the *Arabidopsis* DEMETER and poplar PtDML6 (Fig. 1). Based on this finding, the chestnut protein was named CsDML *(Castanea sativa* DEMETER-like). Biochemical and structural analyses have shown that *Arabidopsis* DEMETER (AtDME), DEMETER LIKE 2/3 (AtDML2, AtDML3) and REPRESSOR OF SILENCING 1 (AtROS1), all grouping in this cluster, are DNA glycosylases that specifically remove 5mC from DNA (Zhu, 2009). The identification of CsDML in this cluster led us to look for the presence of a conserved 5mC DNA glycosylase active site in this protein.

**Figure 1.**
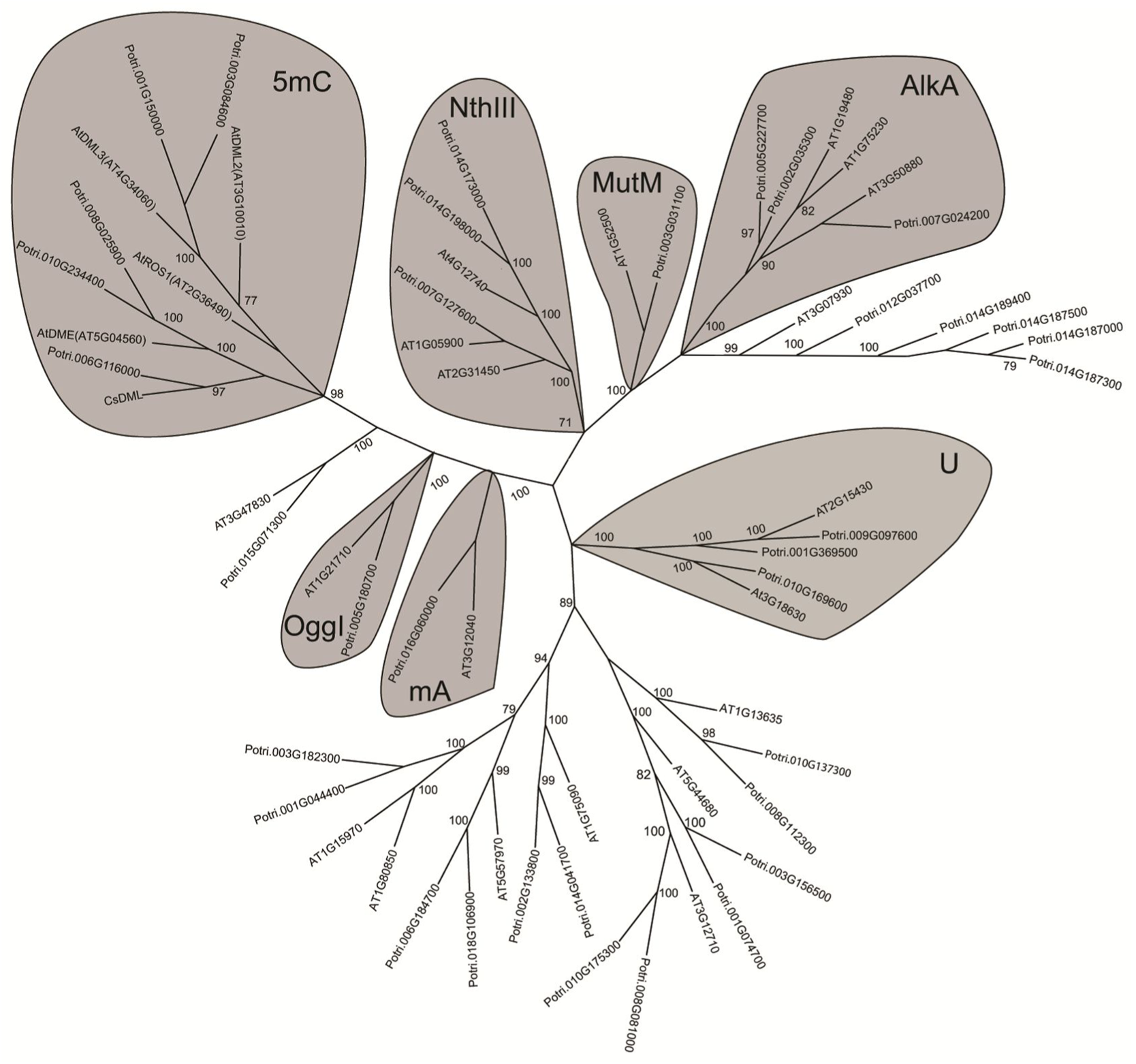
Phylogenetic tree analyses of glycosylase proteins from Arabidopsis, poplar and chestnut. Maximum likelihood tree obtained using the RAxML tool with 100 bootstrap replicates. Gray clades grouped protein sequences closer to proteins of known function: AlkA: alkyladenine glycosylases that excise 3-methyladenine (Begley et al., 1999); Nth: endonuclease III and OggI: 8-oxoguanine glycosylase I, which remove oxidative damaged bases from the DNA backbone (Rosenquist et al., 1997); U: specific for the excision of uracil (Córdoba-Cañero et al., 2010); mA: excise 3-methyl adenine (Santerre and Britt, 1994); MutM: formamidopyrimidine-DNA glycosylase (Duclos et al., 2012); 5mC: the group of 5mC DNA glycosylase (Zhu, 2009). CsDML and 5 poplar DML proteins clustered with AtDME, AtROS1, AtDML2 and AtDML3. Only bootstraps higher than 70 are shown.

### Comparative analyses of the 5-methyl cytosine DNA glycosylase domain suggests a functional catalytic site for CsDML and PtDML6

A MAFFT protein alignment was carried out using the 5mC DNA glycosylases sequences used to generate the phylogenetic tree. The alignment revealed that CsDML, PtDML6, PtDML8 and PtDML10 protein sequences show the three characteristic domains of AtDME/AtDMLs/AtROS1: a lysine-rich domain, a DNA glycosylase domain and a C-terminal domain (Fig. 2A). In AtDME/AtDMLs/AtROS1, the catalytic DNA glycosylase domain consists of two conserved non-contiguous segments connected through a linker region which is highly variable in sequence and length (Ponferrada-Marín et al., 2010). These segments are highly conserved in CsDML, PtDML6, PtDML8 and PtDML10, but partially conserved in PtDML1 and PtDML3 (Fig. 2, A and B).

**Figure 2.**
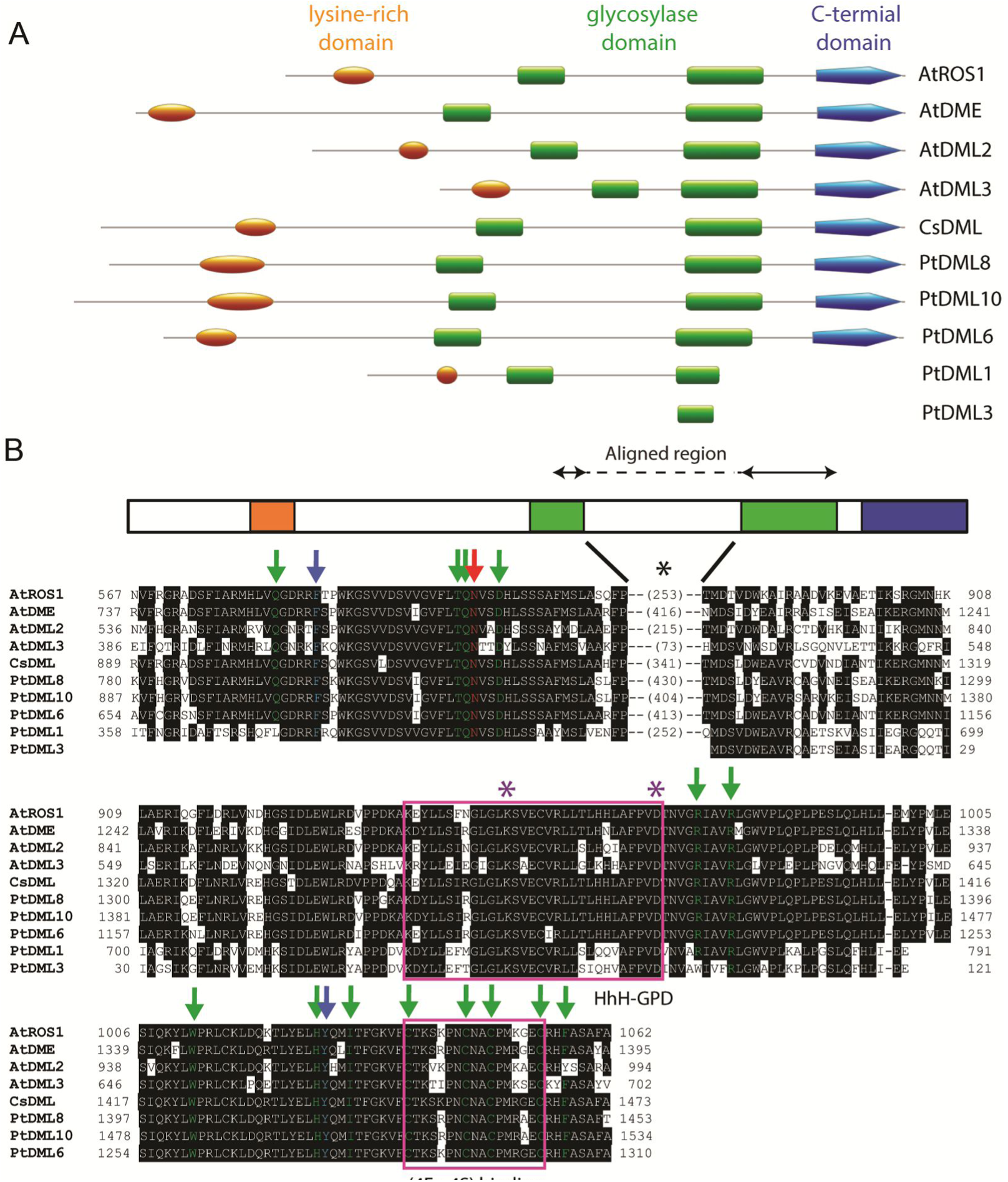
Comparative analysis of the 5mC DNA glycosylase domains of Arabidopsis, poplar and chestnut DMLs. A, Scheme showing the 5mC DNA glycosylase conserved domains in Arabidopsis, poplar and chestnut. CsDML, PtDML6, PtDML8 and PtDML10 conserve all the domains required for 5mC DNA glycosylase activity. Orange oval represents the lysine-rich domain. Green rectangles represent the motifs of the DNA glycosylase domain. Blue pentagon represents the C-terminal domain. B, MAFF alignment of the 5mC DNA glycosylase domain of Arabidopsis, poplar and chestnut. Black boxes indicate identical aminoacids. Black asterisk indicates the non-conserved linker sequence between the two motifs of the DNA glycosylase domain. Arrows indicate the position of the aminoacids necessary for the activity of AtROS1/AtDME/AtDML2-3 (green: required for base 5mC excision, blue: required for preference for 5mC over T mismatches, red: involved in control of context preference for 5mC). Purple boxes indicate HhH-GPD and [4Fe– 4S] motifs. Purple asterisk show the position of the lysine residue necessary for bifunctional glycosylase/lyase activity and the conserved aspartic acid of the DNA glycosylase family.

Structural studies and site-directed mutagenesis of AtROS1 and AtDME have identified the amino acids required for excision of 5mC in the second segment of the DNA glycosylase domain (Gong et al., 2002; Mok et al., 2010; Ponferrada-Marín et al., 2011; Parrilla-Doblas et al., 2013; Brooks et al., 2014). This domain includes the HhH-GPD motif, the invariant aspartate and the iron sulfur [4Fe-4S] cluster loop motif (Fig. 2B). CsDML, PtDML6, PtDML8 and PtDML10 contain all the residues required for the excision of 5mC, suggesting they have a functional catalytic site.

### SD and cold temperatures promote *CsDML* and *PtDML6* expression

Day length shortening and temperature decrease in autumn are the environmental cues driving seasonal growth cessation and bud development in trees (Cooke et al., 2012). Given the seasonal expression of *CsDML,* preferentially at the beginning of autumn, we analyzed the expression of this gene under each inductive condition, separately (SD and cold temperature). By qRT-PCR, we determined that *CsDML* increased its expression in response to both SD and cold temperature signals (Fig. S2, B and C). qRT-PCR analysis also indicated that *CsDML* mRNA expression is not required for dormancy maintenance, since its expression decreased in dormant plants that have not fulfilled the chilling requirement, when these plants were transferred to favorable growth conditions (Fig. S2D). Likewise, qRT-PCR experiments showed that expression of *PtaDML6* increased in response to SD and cold temperature in poplar stems (Fig. S2, F and G). We also assessed the SD-induced expression of *PtaDML6* by fluorescence *in situ* hybridization (FISH) on sections of poplar apical buds. After 24 days under SD, *PtaDML6* mRNA was detected in the shoot apical meristem (SAM) and leaf primordial but not in LD conditions (Fig. 3, A and B). This result showed that the accumulation of *PtaDML6* mRNA in the shoot apex is SD dependent. Taken together, the phylogenetic analysis and SD gene expression data indicate that both *CsDML* and *PtDML6* are induced by environmental conditions that lead to growth cessation and dormancy, and may play a role in the regulation of bud set.

**Figure 3.**
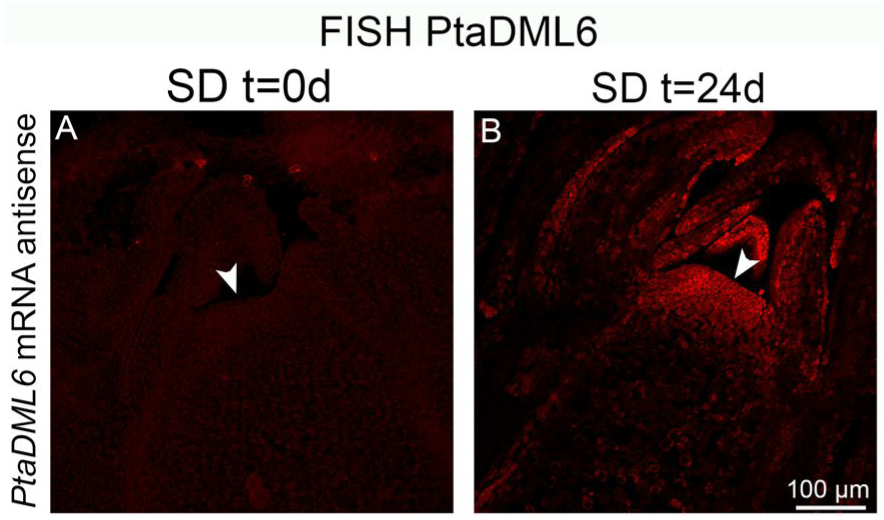
*PtaDML6* fluorescence *in situ* hybridization. *PtaDML6* mRNA detection in 20 μm sections of paraffin-wax embedded WT apical buds at SD time points 0 and 24 days (n=3). The highest intensity of fluorescence was observed in cytoplasmic areas of cells in the meristem and leaf primordia at 24 days under SD conditions. Arrows point at the SAM.

### Apical bud maturation is accelerated in *CsDML* OX poplars

The role of *CsDML* during dormancy was further investigated by overexpressing it in the model species poplar. *CsDML* OX plants showed no phenotypic variation compared to WT when grown under LD conditions. Given their higher expression levels (Fig. S3), *CsDML* OX1 and OX3 lines were chosen to examine growth and apical bud phenology in growth chambers, under photoperiodic and temperature conditions that mimic seasonal transitions from autumn-winter-spring (Rohde et al., 2011b). Phenology of WT and *CsDML* OX1 and OX3 lines did not show any differences until they reached stage 1, once leaf emergence from the apex ceased and green bud scales were formed in response to SD. Then, apical buds from both *CsDML* OX1 and OX3 lines underwent maturation more rapidly, reaching stage 0 earlier than WT buds (Fig. 4A). A deviation in the bud maturation rate was observed from the 30^th^ day of SD exposition onwards. Thus, while WT took 30 days more to reach stage 0, *CsDML* OX1 and OX3 needed 19 and 23 days, respectively (P<0.01) (Fig. 4A). Accordingly, bud scales in *CsDML* OX1 and OX3 plants acquired the red coloration observed normally during bud set earlier than in WT plants (Fig. S3). These results indicate that constitutive expression of *CsDML* in poplar specifically accelerates the transition from stage 1 to 0 during bud set.

**Figure 4.**
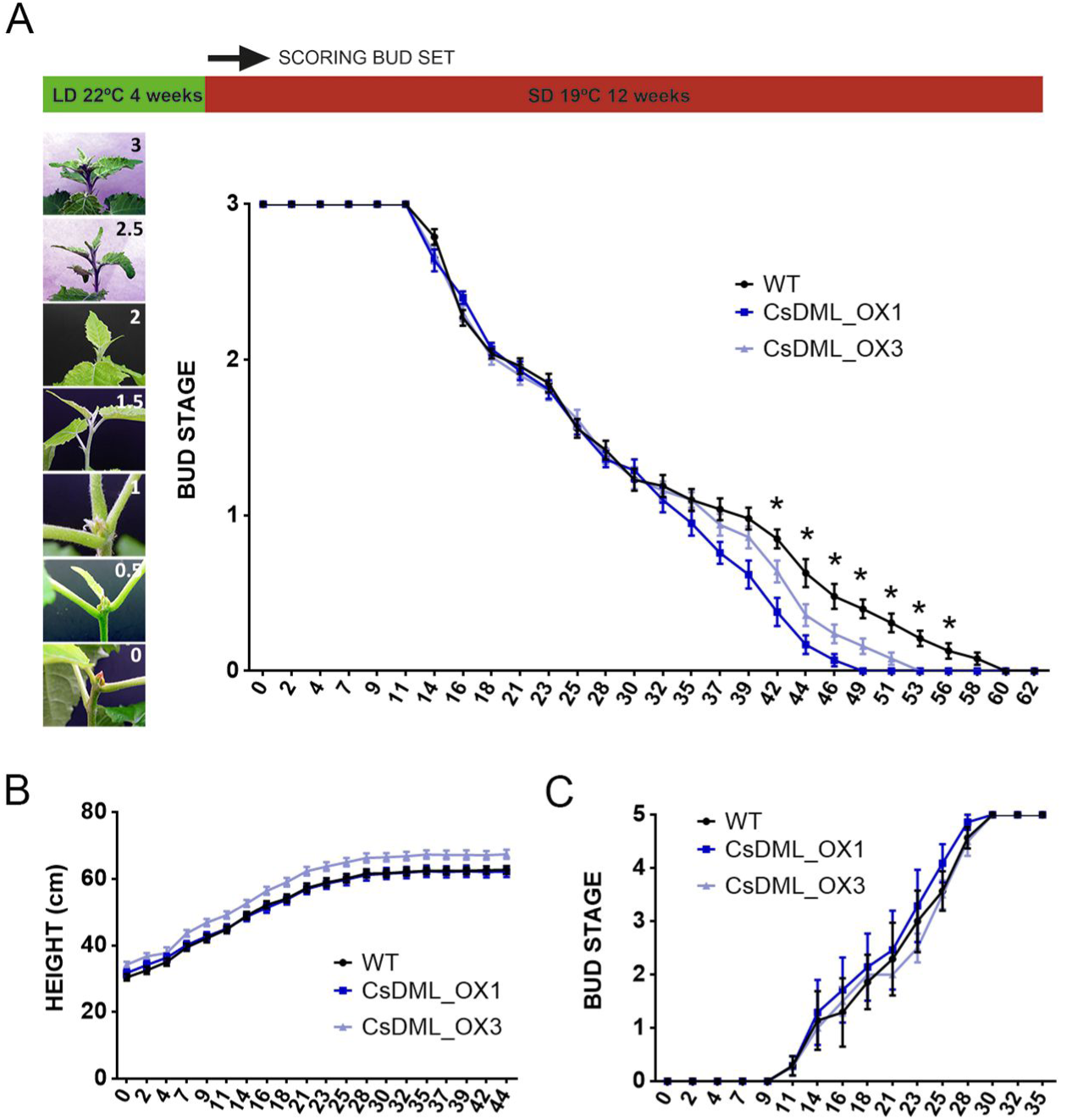
Phenologycal studies of *CsDML* OX plants. *A*, Temporal dynamics of bud stage during the induction of apical bud formation by SD. WT, *CsDML* OX1 and OX3 plants were grown under LD, 22°C conditions for 4 weeks and then the buds were scored over the following 13 weeks after transferring the plants to SD, 19°C conditions. Mean bud scores ±SE are indicated for WT (*n*=24), *CsDML* OX1 (*n*=21) and *CsDML* OX3 (*n*=25). Significant differences between *CsDML* OX1 and WT at t=49 d of SD and between *CsDML* OX3 and WT at t=53 d of SD were analyzed by the Wilcoxon Rank Sum test in R, ***: P<0.01. B, Growth cessation in response to SD. Heights of WT, *CsDML* OX1 and OX3 plants were measured under SD conditions for 44 days. Height means (cm) ±SE are indicated for WT (*n*=24), *CsDML* OX1 (*n*=21) and *CsDML* OX3 (*n*=25). C, Timing of bud burst after dormancy. WT, *CsDML* OX1 and *CsDML* OX3 lines were treated as shown in A, following the shift from SD, 4°C to LD, 22°C, apical growth reactivation was monitored. Mean bud burst scores ±SE are indicated for WT (*n*=7), *CsDML* OX1 (*n*= 7) and *CsDML* OX3 (*n*=7).

Interestingly, an experiment in which we sampled WT, *CsDML* OX1 and OX3 plants at time points t=0, t=24 and t=50 days under SD conditions, all plants arrested their growth on the 28^th^ day of SD exposition (Fig. 4B). Thus, while bud set is affected by *CsDML* overexpression, the cessation of growth elongation is not. Furthermore, once the chilling requirement had been fulfilled, WT, *CsDML* OX1 and OX3 plants were placed under LD, 21°C conditions and bud burst was monitored. After 18 days, all three genotypes displayed green buds (stage 1 to 2), and after 30 days all three genotypes had elongated their shoots (stages 3 to 5) (Fig. 4C). These results suggest that the constitutive expression of *CsDML* does not contribute to bud burst.

### 5mC immunodetection studies suggest a locus-specific role for DNA demethylase during bud maturation

Overall changes in 5mC levels were compared in poplar apex sections of WT, *CsDML* OX1 and OX3 plants during SD induction of bud formation in growth chamber. By immunolocalization we examined overall DNA methylation levels under LD conditions (t=0) and after 24 days of growth under SD conditions (t=24), that is, six days before the apices of *CsDML* OX lines began to show faster apical bud maturation than WT plants (Fig. 5, A-F). This experiment revealed that at t=0, DNA methylation levels are restricted to the central zone of the SAM (Paul et al., 2014) both in WT and *CsDML* OX apices (Fig. 5, A-C). At t=24, DNA methylation spread to the rest of the SAM and other bud regions (Paul et al., 2014), with the same pattern in WT and *CsDML* OX apices (Fig. 5, D-F). Quantification of the fluorescence derived from DNA methylation indicated that the ratio of levels in the central zone of the meristem and the remaining SAM was stable in the different genotypes at both time points (Fig. 5, G-I). All these findings indicate *CsDML* overexpression does not affect the overall DNA methylation changes induced by SD, thus suggesting a locus-specific activity for this gene during bud development.

**Figure 5.**
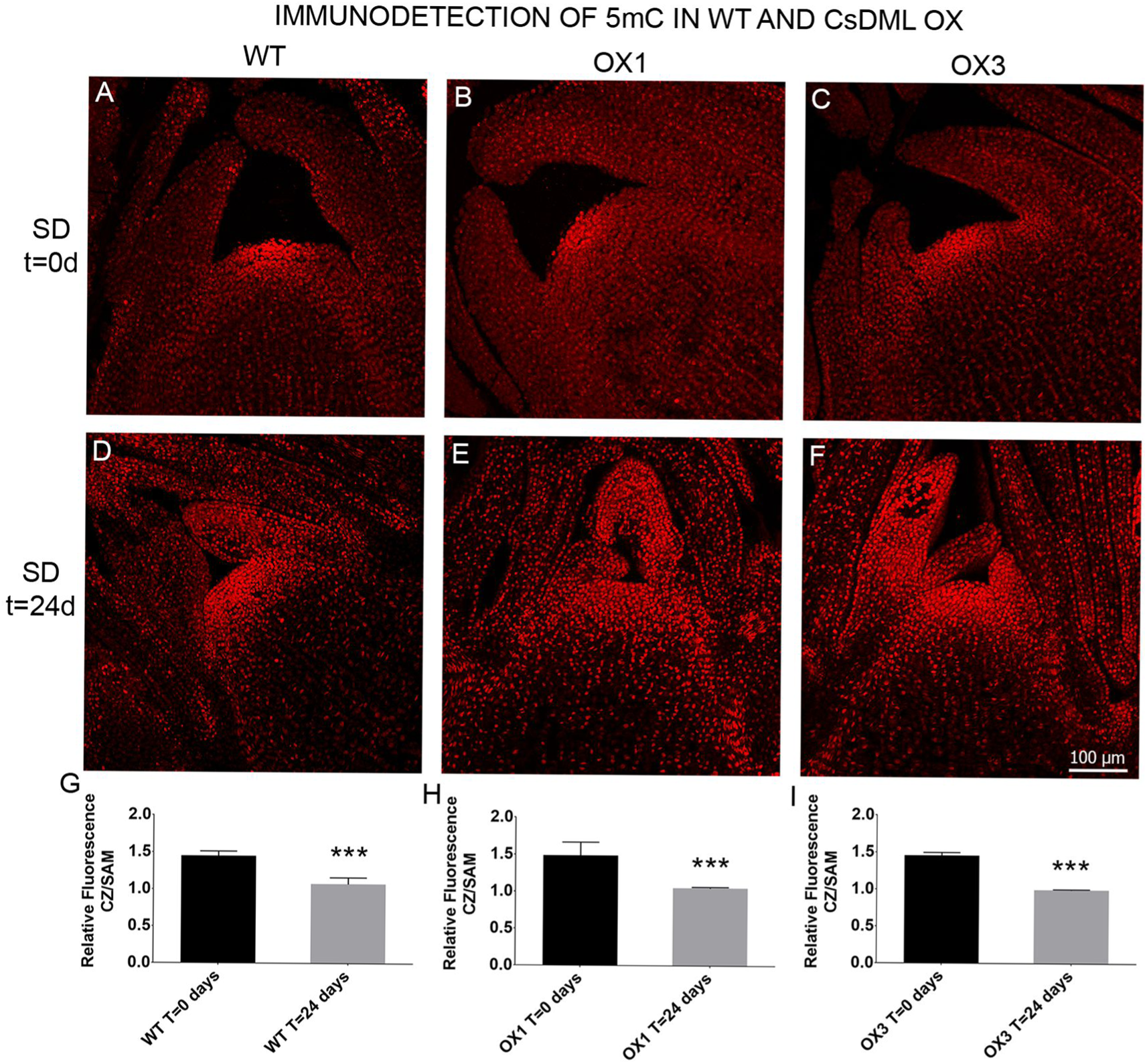
Immunodetection of 5mC in *CsDML* OX and WT. Immunofluorescence detection of 5-methylcitidine in 20 μm sections of paraffin-embedded poplar apical buds obtained from the WT (*n*=3) (A,D) and lines *CsDML* OX1 (*n*=3) (b,e), *CsDML* OX3 (*n*=3) (C,F) (4-week-old trees). Apical bud showing the 5mC signal both in LD conditions (t=0) (A,B,C) and after 24 days of growth under SD conditions (t=24 d) (d, e, f). Histogram represents the ratio between fluorescence signals emitted by the central zone (CZ) and the whole SAM of WT (G), *CsDML* OX1 (H), *CsDML* OX3 (I) at t=0 vs. at t=24 d (means ±SE). Differences between each time point and genotype were assessed by one-way ANOVA. *** indicates significant differences between the ratio t=24 d to t=0 in each genotype (P<0.01).

### *CsDML* OX plants differentially express genes involved in bud formation

To investigate the molecular basis causing the acceleration of bud maturation in *CsDML* OX plants, a genome wide transcriptome analysis was performed by RNA sequencing. WT and *CsDML* OX1 and OX3 plants were first grown in LD conditions during 4 weeks and then in SD conditions for additional 4 weeks. Apices were collected on day 28, just before the early bud maturation took place in *CsDML* OX plants (Fig. 4A). Total RNA was extracted from 25 pooled apices per genotype and RNA-seq libraries were generated and sequenced. Transcriptome comparisons identified 318 DEG in *CsDML* OX lines compared to the WT, among which 79 were found to be upregulated and 239 downregulated (Table S1). To examine their molecular functions, GO analysis of these DEGs was conducted (Table S2). Upregulated genes in *CsDML* OX lines did not show significant enrichment in any biological categories. Downregulated genes showed enrichment in the following categories: cellular processes, cell cycle and cell division, DNA and chromatin modifications, anatomical structure development, and developmental processes (Table S2).

Upregulation of flavonoid biosynthesis have been shown as molecular markers of bud maturation (Ruttink et al., 2007; Tylewicz et al., 2015). Our transcriptome analyses revealed three upregulated genes in *CsDML* OX plants encoding putative enzymes belonging to the flavonoids biosynthetic pathway. They were a chalcone synthase (CHS), a quercetin 3-O-glucosyltransferase (3GT) and a dihydroflavonol 4-reductase (DFR) (Table S1). CHS and DFR enzymes are known to be involved in the early steps of flavonoid biosynthesis (Ono et al., 2010), so upregulation of these genes in *CsDML* OX plants could correlates with their early red coloring during bud maturation (Perry, 1971; Tylewicz et al., 2015).

The induction of *PtaCHS, PtaDFR* and *Pta3GT* mRNAs, codifying for enzymes belonging to the flavonoids biosynthetic pathway, were confirmed by qRT-PCR studies, showing their differential accumulation after 21 days under SD conditions in apices of *CsDML* OX plants compared to WT plants. Expression of *PtaCHS* was 2.7 and 2.3 times higher in *CsDML* OX1 and *CsDML* OX3 than in WT. *PtaDFR* was 9 and 4.6 higher in *CsDML* OX1 and *CsDML* OX3 than in WT and *Pta3GT* was 3.7 and 2.1 higher in *CsDML* OX1 and *CsDML* OX3 than in WT (Fig. 6A).

**Figure 6.**
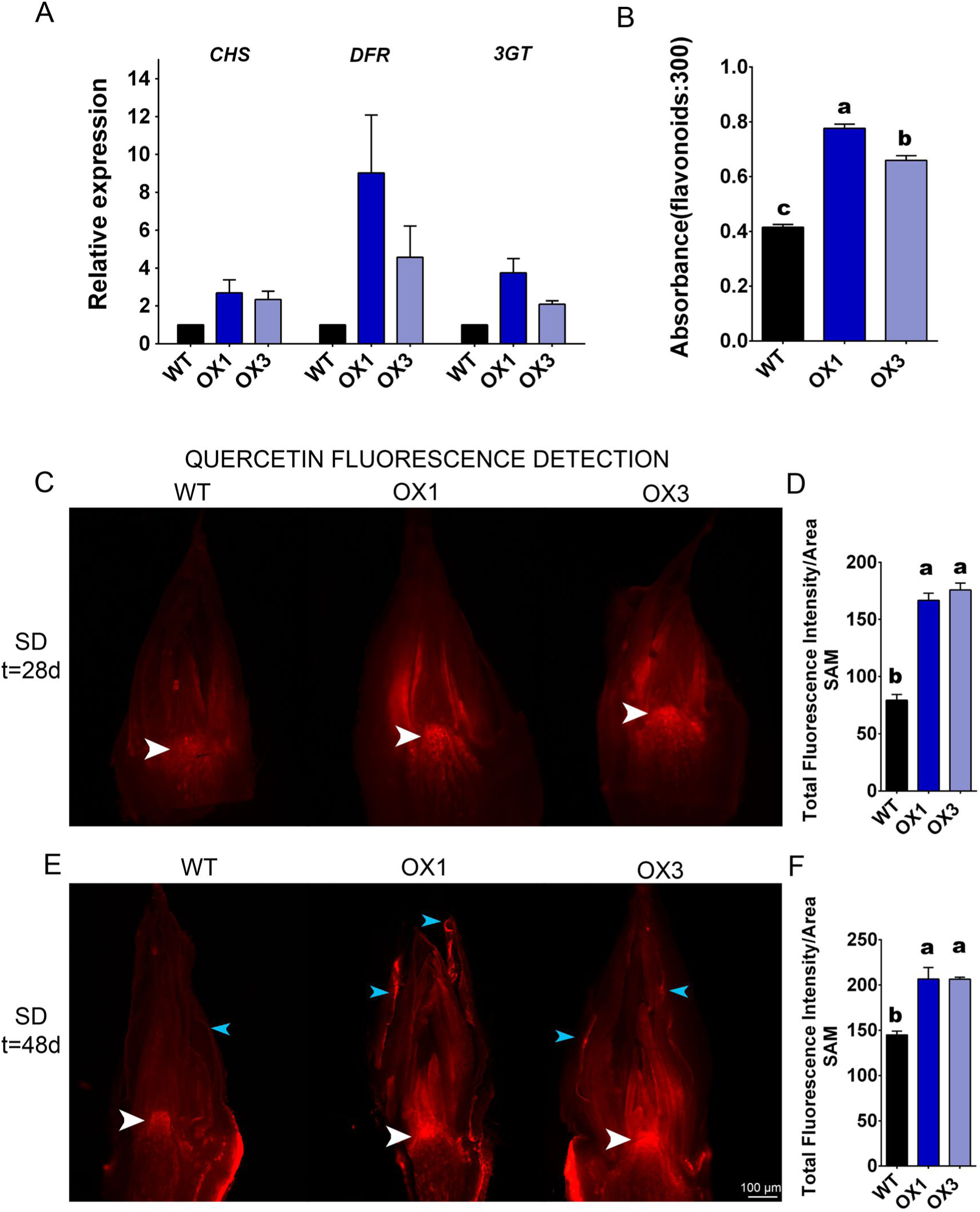
Flavonoids accumulation assays in *CsDML* OX and WT plants apices. A, Quantitative RT-PCR analysis of *PtaCHS, PtaDFR* and *Pta3GT* in *CsDML* OX1, *CsDML* OX3 and WT apices after 21 days in SD. Plotted values and error bars are fold-change means ± SE recorded in two biological replicates. B, Absorbance measurements of flavonoid (at 300nm) in *CsDML* OX1, *CsDML* OX3 and WT apex extracts after 48 days in SD. Values are the means of three biological replicates, and bars indicate SE. Significant differences between genotypes were analyzed by ANOVA and the TUKEY test. Different letters showed significant differences (P<0.05). C, Localization of Quercetin-DPBA fluorescence emission in *CsDML* OX1, *CsDML* OX3 and WT apices after 28 days in SD. White arrows indicate the SAM areas. D, Quantification of Quercetin-DPBA fluorescence in the SAM in *CsDML* OX1, *CsDML* OX3 and WT apices after 28 days in SD. Values are the means of three independent samples, and bars indicate SE. Significant differences in genotype were analyzed by ANOVA and the TUKEY test. Different letters showed significant differences (P<0.05). E, Localization of Quercetin-DPBA fluorescence emission in *CsDML* OX1, *CsDML* OX3 and WT apices after 48 days in SD. White arrows indicates the SAM areas and blue arrows indicate the external bud scales. F, Quantification of Quercetin-DPBA fluorescence in the SAM in *CsDML* OX1, *CsDML* OX3 and WT apices after 48 days in SD. Values are the means of three independent samples, and bars indicate SE. Significant differences between genotypes were analyzed by ANOVA and the TUKEY test. Different letters showed significant differences (P<0.05).

### *CsDML* OX accumulates flavonoids earlier than WT in SAM and bud scales

To determine whether an increment of flavonoid biosynthetic genes correlates with flavonoid accumulation, we quantified the absorbance of flavonoids in apical buds of WT, *CsDML* OX1 and *CsDML* OX3 after 48 days in SD. Both transgenic lines, showed an increment of the absorbance of flavonoids compared to WT, indicating the involvement of CsDML in the early accumulation of flavonoids (Fig. 6B).

The spatio-temporal accumulation of flavonoids was explored by detecting quercetin, a flavonoid of the flavonol subgroup that can be identified by fluorescence after staining with DPBA. We localized and quantified quercetin in apical buds of *CsDML* OX1, *CsDML* OX3 and WT plants, after 28 and 48 days of SD induction. Apical buds were already formed after 28 days under SD conditions, although still immature, in WT, *CsDML* OX1 and *CsDML* OX3 trees. At this time point, the red fluorescence emitted by quercetin-DPBA is primarily expressed in SAM. However, higher fluorescence intensity was detected in *CsDML* OX1 and *CsDML* OX3 SAM, compared to WT (Fig. 6, C and D). After 48 days, bud maturation occurred earlier in the OX than in WT plants, as indicated by the more intense red-brown coloration (Fig. 4 and S3). In the SAM of the three genotypes, fluorescence was higher after 48 days than after 28 days, suggesting a more significant accumulation of quercetin in SAM under SD (Fig. 6, C-F). Accordingly, fluorescence intensity was higher in the SAM of the transgenic lines than in WT. After 48 days in SD, quercetin-DPBA fluorescence is also highly detected in external bud scales of both transgenic lines (Fig. 6, E and F). The localization of quercetin after SD induction matched with the *PtaDML6* spatial expression pattern, suggesting a conserved function between CsDML and PtaDML6, in promoting flavonoid biosynthesis.

Altogether, these results showed that *CsDML* promotes the induction of flavonoid biosynthesis enzymes, needed for apical bud maturation through the accumulation of flavonoids in the SAM and bud scales, during SD induced bud formation.

## DISCUSSION

### A putative locus-specific 5mC DNA demethylase is involved in bud maturation

Here we show that *CsDML* is preferentially expressed at the beginning of autumn in response to SD and cold temperatures. Our qRT-PCR and RNA FISH studies confirm the induction by SD of the poplar closest orthologue of the chestnut *CsDML, PtDML6,* in the apical meristem. Previous RNA-seq studies have associated *PtDML6,* with dormancy induction, in poplar stem and apex (Shim et al., 2014; Fig. S4). These results suggest that activation of a 5mC demethylase during bud maturation is conserved both in chestnut and poplar. SD treatment promotes an overall increase in 5mC DNA methylation in SAM and leaf primordial tissues. Such increased levels of 5mC DNA methylation have been previously observed during winter in chestnut and poplar (Santamaría et al., 2009; Conde et al., 2013). Also, the spatial pattern of the *PtDML6* transcript in apical buds detected by FISH overlaps with that of 5mC immunofluorescence methylation observed under SD-induction conditions. Overall 5mC levels in apical buds were unaffected by *CsDML* overexpression in poplar, both under LD conditions and following 24 days of exposure to SD. Other authors have reported a locus-specific 5mC DNA demethylation in response to stress for *Arabidopsis* ROS1, DML2 and DML3 (Zhu et al., 2007; Yu et al., 2013; Le et al., 2014). On the basis of these results, we propose that PtDML6 and CsDML act as locus-specific 5mC DNA demethylases during bud maturation.

### CsDML accelerates apical bud maturation

During SD-induced bud development, two concatenated events can be identified: cessation of shoot elongation and apical leaf formation, and the progressive arrest of cell proliferation while the buds are formed (Rohde et al., 2002). The later includes differentiation of leaf primordia into bud scales alongside with the formation of the embryonic shoot. Bud scales accumulate flavonoid components leading to their characteristic red-brown color (Rohde et al., 2002; Tylewicz et al., 2015). In addition to the above-mentioned transformations, the SD signal promotes the acquisition of cold and desiccation tolerance in apical and cambial meristems before they become dormant (Rohde et al., 2002; Rohde and Bhalerao, 2007; Ruttink et al., 2007; Cooke et al., 2012).

So far, published functional data for the regulation of bud formation were obtained by modifying ethylene and abscisic acid (ABA) signaling pathways (Rohde et al., 2002; Ruonala et al., 2006). Phenological studies on ethylene-insensitive birch trees *(Betula pendula)* showed that these trees do not form apical buds under inductive conditions, indicating that ethylene signaling is essential for bud formation but not for the cessation of shoot elongation (Ruonala et al., 2006). In poplar, downregulation of *ABSCISIC ACID INSENSITIVE 3 (ABI3)* or its interactor *FDL1,* promotes early bud formation without affecting the cessation of shoot elongation (Rohde et al., 2002; Tylewicz et al., 2015). Anatomical analysis of *PtABI3* overexpressing and knockdown plants have revealed deficiencies in size and ratio of embryonic leaves and bud scales under SD conditions, pointing to a role of ABA in these processes (Rohde et al., 2002). Our observations suggest that overexpression of *CsDML* only impacts the formation of the bud, while growth arrest continues unaltered. This suggests that the activity of the CsDML protein requires other SD dependent factors (or that must first be removed) for DML to function. In our transcriptional profiling of *CsDML* OX compared to WT, we did not detect enrichment either for ABA or for ethylene pathway enzymes, even though several ABA-related genes such as *LATE EMBRYOGENESIS ABUNDANT (LEA)* were found among the DEG (Table S1). Therefore, we propose that our 5mC DNA demethylase controls bud formation separately from ethylene and may act downstream or independently of ABI3. Because *CsDML* OX plants did not differ phenotypically from WT plants until the 4th week of SD treatment, CsDML may act jointly with the ABA signal, which accumulates in apical buds at the same time after SD induction (Ruttink et al., 2007).

### Overexpression of CsDML promotes flavonoid biosynthesis

Phenylpropanoids pathway has been extensively investigated in model tree species due to the increasing interest of renewable energy resource (Li et al., 2014). The transition from active growth to dormancy has been associated to a decreased cellulose biosynthesis, concomitant to the generation of phenylpropanoids (lignin and flavonoids) (Perry, 1971; Rohde et al., 2002; Tylewicz et al., 2015). Accordingly, our comparative RNA-seq analyses of SD-induced apical buds revealed three upregulated flavonoid biosynthesis genes and two downregulated cellulose synthase-like genes in *CsDML* OX plants compared to WT. By qRT-PCR we confirmed that *CHS, DFR* and *3GT* expression was upregulated in transgenic plants, and the measurement of flavonoids absorbance revealed higher levels of these compounds in transgenic plants apices, concurrent with the earlier accumulation of red color in apical buds that was observed in *CsDML* OX earlier than in WT plants. Flavonoids accumulation as part of the adaptive response and bud maturation changes, occur in response to seasonal dormancy (Ruttink et al., 2007; Tylewicz et al., 2015). These compounds could act as a sun screen protecting agents against photooxidative damage of the organs present in buds (bud scales, embryonic leafs, leaf primordia and SAM) (Falcone Ferreyra et al., 2012). CHS and DFR are involved in the first steps of the flavonoid biosynthesis pathway implying high levels of synthesis of anthocyanins, flavonols and flavanols (Ono et al., 2010), suggesting that DML accelerated the accumulation of flavonoids during apical bud formation. The regulation of the flavonoid pathway through DNA methylation has been recently reported in tobacco (Bharti et al., 2015). Tobacco plants overexpressing *AtROS1* showed a higher expression of flavonoid biosynthesis enzymes, including CHS, CHI and DFR, and lower levels of 5mC in their promoters than in WT plants (Bharti et al., 2015). This scenario was enhanced under salt stress, suggesting that a stimulation of AtROS1 activity under these conditions activates flavonoid pathway genes (Bharti et al., 2015). The increased levels of flavonoid biosynthetic enzymes detected in tobacco *AtROS1* and poplar *CsDML* overexpressors indicated that this pathway could be activated by 5mC demethylation in both species.

Downregulation of cell proliferation genes precedes meristem inactivation at the final stage of bud development (Ruttink et al., 2007). Several cell cycle regulators were downregulated in *CsDML* OX plants compared to the WT when they were exposed to 28 days of SD. A pivotal role of flavonoids in the cell cycle arrest has been reported in both plants and animals (Taylor and Grotewold, 2005; Woo et al., 2005; Butelli et al., 2008). The build-up of flavonoids, such as quercetin and other molecules synthesized in the first steps of the flavonoid biosynthesis pathway, is known to block polar auxin transport, required for cell division and differentiation in SAM tissues (Kuhn et al., 2011). According to this, the flavonol quercetin was found earlier accumulated in the SAM and bud scales of *CsDML* OX than in WT plants. The accumulation of quercetin in SAM tissues undergoing cell arrest could be a consequence of auxin transport blockage triggered by flavonoid. In future studies, the downregulation of cell proliferation genes during bud formation should be explored as a consequence of the build-up of flavonoids.

## CONCLUSION

Based on our observations, we propose the following model for the implication of a DML in the SD-mediated control of bud maturation. As the day-length shortens, *DML* mRNA accumulates in poplar SAM and leaf primordial. DML acts as a locus-specific 5mC DNA demethylase, activating genes encoding enzymes of the flavonoids biosynthesis pathway. These flavonoids will be implicated in bud maturation through: the acquisition of the characteristic red-brown color, acting as a sunscreen protecting the organs enclosed within the buds against photo oxidative damage and the repression of cell proliferation, which ultimately could be triggering the acceleration of bud formation. This study provides *in planta* evidence for a role of 5mC DNA demethylase activity in the regulation of bud maturation through the biosynthesis of flavonoids.

## MATERIAL AND METHODS

### Plant materials, growth conditions and assays

To monitor gene expression throughout the year, 2-year-old branches were harvested monthly from adult European chestnut trees *(Castanea sativa* Mill.) growing under natural conditions in Zarzalejo (Madrid, Spain 4°11´W, 40°35´N). At each time point, samples were collected from three different chestnut tree individuals. To evaluate gene expression induced by short day (SD) conditions, plants were first grown under long day (LD) (16h light/8h darkness) at 22°C during 8 months, and then changed to SD (8h light/16h darkness) at 22°C for 6 weeks. In each time point, stems of 4 plants were sampled and pooled. To evaluate gene expression induction by low temperatures (4°C), plants were grown under LD conditions at 22°C during 8 months and then changed to LD and 4°C during 7 weeks. In each time point, stems of 4 plants were sampled and pooled. For the dormancy release experiment, 11-month-old dormant chestnut plantlets grown under natural conditions were transferred to a growth chamber under LD and 22°C before the chilling requirement was fulfilled. Stems from 3 plantlets were sampled and pooled at the indicated time points. Two biological replicates were analyzed. The details of these experiments have been described elsewhere (Ramos et al., 2005).

To evaluate gene expression induction by SD in poplar *(Populus tremula x alba* INRA clone 717 1B4), plants were first grown under LD and 22°C during 6 weeks, and then changed to SD and 22°C during 7 weeks. In each time point, stems of 2 plants were sampled and pooled. To evaluate the induction by cold temperatures in poplar, plants were first grown under LD, 22°C conditions during 8 weeks and then changed to 4°C during 7 weeks. In each time point, stems of 3 plants were sampled and pooled.

For the phenological assays, *in vitro*-cultivated poplar plantlets *(P. tremula x alba* INRA clone 717-1B4) of wild type (WT) and two selected independent CsDML-overexpessing lines, *CsDML* OX1 and OX3, were transferred to 3.5 L pots containing blond peat, pH 4.5 (Moreno-Cortés et al., 2012). Before any photoperiodic treatments, plants were grown in a chamber under controlled conditions: LD, 22°C, 65% relative humidity and 150 μmol m^-2^ s^-1^ photosynthetic photon flux). After 4 weeks these plants were exposed to a SD, 19°C regime for 10 weeks to induce bud set. Bud set progression was graded by scoring from stage 3 (fully growing apex) to stage 0 (fully formed apical bud) according to Rohde et al. (2011b). Apical buds for immunofluorescence were sampled at 0 and 24 days after SD induction; buds used for transcriptome analysis were sampled 28 days after SD induction; apical buds for quercetin fluorescence detection were sampled at 28 and 48 days after SD induction and apical buds for flavonoids absorbance measurements by spectrophotometry were sampled after 48 days in SD, in three independent assays. To break dormancy and allow further bud burst under good conditions (LD, 22°C), poplars plants were subjected to a SD, 4°C regime for 4 weeks. The regrowth was scored according to the six developmental stages of bud burst (stage 0 to stage 5) according to UPOV (International Union for the Protection of New Varieties of Plants, 1981) and Ibáñez et al. (2010). Throughout the entire experiment, photosynthetic photon flux and relative humidity were kept at 150 μmol m^-2^ s^-1^ and 65%, respectively. Statistical significance on differences in the phenological apex stages between *CsDML* OX and WT plants was tested through pairwise comparisons using the Wilcoxon Rank Sum test after Shapiro-Wilk confirmation of the non-normal distribution of data. All statistical tests were run in R (www.r-project.org).

To evaluate the expression levels of *PtaCHS, PtaDFR* and *Pta3GT* in transgenic and WT poplar plants, *CsDML* OX1, *CsDML* OX3 and WT in 3 weeks old plants were grown under SD conditions during 3 weeks. A pull of 3 apical buds was taken at this time point to analyze gene expression in each sample. Two biological replicates, three technical replicas for each one, were performed.

### Isolation of full-length *CsDML* cDNA

A 464-bp-long fragment of chestnut *DEMETER-like (CsDML)* cDNA was obtained from a winter-enriched cDNA subtractive library (PCR-Select cDNA Subtraction Kit, Clontech Laboratories, Mountain View, CA, USA) prepared from 2-year-old branches of adult trees (Ramos et al., 2005). Full-length cDNA was obtained using the SMARTer RACE cDNA amplification kit (Clontech Laboratories). *CsDML* nucleotide sequence was deposited in the GenBank sequence databases with the accession number KX575710.

### Phylogenetic analysis of HhH glycosylase family members and in-silico identification of the active site for DNA demethylation in the CsDML protein

Sequences of proteins containing the hairpin-helix-hairpin (HhH) glycosylase domain, a characteristic feature of the CsDML protein, were retrieved from *Arabidopsis thaliana* genome using the TAIR10sd website (www.arabidopsis.org). These sequences were used as queries to BLAST the *Populus trichocarpa* proteome (Phytozome v10.3, www.phytozome.net) and identify homologue proteins in this organism. All the protein sequences retrieved from *Arabidopsis* and poplar, together with CsDML, were aligned using MAFFT (E-INS-i algorithm) (Katoh and Standley, 2013) and edited in Jalview 2.8 (Waterhouse et al., 2009). The best evolutionary model of the alignment was highlighted using ProtTest 2.4 (Abascal et al., 2005). The maximum likelihood tree was inferred with the RAxML tool (Stamatakis, 2006) and visualized in the iTOL website http://itol.embl.de (Letunic and Bork, 2007). Bootstrap values were generated with 100 replicates. Putative members of the 5mC DNA glycosylase family were named PtDML, followed by a number corresponding to the chromosome where they are located.

An additional alignment using MAFFT (E-INS-i algorithm) (Katoh and Standley, 2013) was performed with 5mC DNA glycosylases of *Arabidopsis thaliana* (AtDMLs), *Populus trichocarpa* (PtDMLs) and *Castanea sativa* (CsDML). The result was visualized in Jalview 2.8 (Waterhouse et al., 2009). The conserved regions of these proteins were assembled in the PROSITE website (http://prosite.expasy.org/mydomains/). allowing identification of the catalytic center residues in the CsDML protein.

### Quantitative RT-PCR expression analysis

The procedures used for total RNA extraction, synthesis of single-stranded cDNA, design of gene-specific primers, quantitative real-time PCR (qRT-PCR) and data analysis have been previously described in Ramos et al. (2005)and Ibañez et al. (2008). 18S was previously used as reference gene (Ramos et al., 2005; Ibañez et al., 2008; Berrocal-Lobo et al., 2011; Hsu et al., 2011) showing to be stably expressed in all the different conditions used for this study both in chestnut and poplar. The primers used are described in Fig. S1A.

### Generating constructs in a binary vector and poplar transformation

The coding sequence (CDS) of the *CsDML* gene was amplified from chestnut cDNA using PfuUltra Hotstart High-Fidelity DNA polymerase (Agilent Technologies, La Jolla, CA, USA) and the primers used are described in Fig. S1B. The *attB*-flanked PCR product was purified, inserted in the pDONR222 vector (Life Technologies, Carlsbad, CA, USA) and verified by sequencing. The insert was transferred into the destination binary vector pGWB15, which includes a cauliflower mosaic virus (CaMV) 35S promoter and adds 3 copies of hemagglutinin (3xHA) at the N-terminal protein end (Nakagawa et al., 2007). The generation and selection of hybrid poplars overexpressing *CsDML,* hereafter referred to as *CsDML* OX lines, was carried out as described in Moreno-Cortés et al. (2012). Based on qRT-PCR analysis of *CsDML* transgene expression in these plants, we selected 2 lines, *CsDML* OX1 and *CsDML* OX3, for further study.

### Immunofluorescence of 5mC in SD-induced shoot apices

Overall DNA methylation in 5mC residues was examined by immunofluorescence in SD-induced (for 24 days) and non-induced shoot apices of *CsDML* OX and WT hybrid poplars *(CsDML* OX1, *CsDML* OX3 and WT). These apices were formaldehyde-fixed and preserved as previously described in Conde et al. (2013)before embedding in paraffin wax. For this, the specimens were washed twice in PBS (30 min each) and dehydrated in a series of 30%, 50%, 70% (1 h each) and 95% ethanol overnight. Specimens were then washed in 100% ethanol for 12 h. Ethanol was then replaced with a mixture 50% histoclear (HC)-50% ethanol overnight and 100% HC for 12 h (twice). Vials containing the apices were filled up to a quarter of their volume with paraplast and incubated at 42°C for 12 h. Next, another quarter of volume was added, followed by incubation at 60°C overnight. Paraplast in HC was replaced with melted paraffin wax (60°C), and this was replenished every 12 h for 5 days. Finally, the apices were placed in molds and, after solidification at room temperature, they were kept at 4°C. Sections of 15-20 μm were obtained in a microtome Leica 2055 (Leica Microsystems, Wetzlar, Germany) and the paraffin was removed in HC for 15 min and then in 100% ethanol for 15 min. The sections were rehydrated in a series of 96% and 70% ethanol for 10 min each and, finally, in PBS for 15 min. Images of immunolocalization were acquired using a Leica TCS-SP8 confocal microscope under the laser excitation line of 561nm. Gain and offset conditions were maintained in all captures for comparison purposes. Signal quantification in apex sections were performed as reported in (Conde et al., 2013). Three independent samples were analyzed for each genotype and time point.

### Probe synthesis and Fluorescence *in situ* hybridization (FISH)

A specific probe to the mRNA of *PtaDML6* was synthesized from a fragment of 1100-pb-long of the 5´of the Untranslated Terminal Region (UTR). The *PtaDML6* DNA was cloned into pGem-T (Promega, Madison, WI, USA). The probe was *in vitro* transcribed to obtain digoxigenin-UTP (DIG) labeled RNA probes using T7 and Sp6 RNA polymerases for the sense and the anti-sense mRNA probes, respectively (Roche Applied Science, Mannheim, Germany). The probe of 1100- pb-long was subjected to a mild carbonate hydrolysis step to obtain aprox. 100-pb-long RNA fragments before hybridization (Cox et al., 1984).

The specimens for *FISH* were collected from WT poplar trees growth under SD at time points 0 and 24 days. Apexes were fixed, paraffin-wax embedded and cut as described before for the immunolocalization of global 5mC. After deparaffination, dehydration and rehydration, as previously described, bud sections were treated with 2% cellulase and proteinase K (1 μg ml^-1^) at 37°C, for 1 hour each. Afterwards, 30 ng μl ^-1^ of either the sense or the antisense probe diluted in the hybridization mixture (50% formamide, 10% dextrane sulfate, 10 mM Pipes, 1 mM EDTA, 300 mM NaCl, 1000 ng μl ^-1^ tRNA) was applied to the sections and incubated at 50°C overnight. After hybridization the slides were washed in 1X Saline Sodium Citrate buffer (SSC) for 30 minutes at 50°C. DIG detection and fluorescence visualization were carried out with a primary anti-DIG (Sigma) antibody, diluted 1/3000 in 3% BSA in PBS for 1 hour at room temperature, and a secondary anti mouse 546 (ALEXA, Molecular Probes) antibody, applied 1/25 in 3% BSA in PBS for 1 hour at room temperature in the dark. Images were acquired using a Leica TCS-SP8 confocal microscope under the laser excitation line of 561 nm. Gain and offset conditions were maintained in all captures for comparison. Three independent samples were analyzed for each genotype and time point.

### Quercetin fluorescence detection

Quercetin accumulation was examined by fluorescence in apical buds of *CsDML* OX1, *CsDML* OX3 and WT plants after 28 and 48 SD, using diphenylboric acid-2-aminoethyl ester (DPBA, Sigma-Aldrich) dye (0.25% in water). Quercetin-DPBA fluorescence was collected on a Leica MZ10F stereomicroscope with a GREEN filter (excitation BP 546/10, emission LP 590). Images were recorded with CCD Leica DFC 420C camera. The specificity of the signal was already tested for this flavonoid (Silva-Navas et al., 2016). Three independent samples were analyzed for each genotype and time point.

### Flavonoids measurements

To determine the total flavonoid content of *CsDML* OX1, *CsDML* OX3 and WT plants after 48 days in SD, 100 mg of fresh weight of a pull of 4 apical buds for each sample, were homogenized in HCL:methanol (1:99, v/v). The tissue was shaken at 4°C overnight. The supernanant absorbance was determined in a Ultraspec 3300 Pro spectrophotometer (Amersham Biosciencies) after diluting to ¼ with HCL:methanol (1:99, v/v) at 300 nm (Nogués and Baker, 2000). Three biological replicates were performed for this experiment for each genotype and time point.

### RNA sequencing in SD-induced shoot apices and identification of differentially expressed transcripts

Expression profiles were examined by RNA sequencing (RNA-Seq) in SD-induced shoot apices of *CsDML* OX and WT hybrid poplars. At 28 day in SD, 25 apices were collected from 25 individuals of WT, *CsDML* OX1 and *CsDML* OX3 trees. Previous to RNA extraction, apices were dissected under a stereomicroscope to remove the leaflets surrounding the shoot apical meristem. RNA extraction was performed following earlier published protocol (Ibañez et al., 2008). Libraries were prepared using 1 μg of RNA in the NEBNext^®^ Ultra™ Directional RNA Library Prep Kit following the supplier's instructions (New England Biolabs, Ipswich, MA, USA). Sequencing was done with an Illumina NextSeq500 in high-throughput mode (2×150 cycles) at the Interdisciplinary Center for Biotechnology Research at the University of Florida (Gainesville, FL, USA). Raw data for the RNAseq was deposited in the Gene Expression Omnibus database at NCBI, with GEO accession number GSE87155 (Edgar et al., 2002; Barrett et al., 2013).

Pre-processing and assembly of sequence data were conducted using the web-based platform Galaxy at https://usegalaxy.org/ (Giardine et al., 2005; Blankenberg et al., 2010; Goecks et al., 2010), which includes tools to remove reads containing N blurs, adapter sequences, and sequences shorter than 15 nucleotides. Quality control of reads was carried out with the FastQC Galaxy tool (Andrews, 2010). Removal of rRNA, if needed, was performed locally with the SortMeRNA tool (Kopylova et al., 2012). The TopHat Galaxy tool was used to map clean reads to the *Populus tremula x P. alba* INRA clone 717-1B4 genome using the default settings (Kim et al., 2013). We used the *summarizeOverlaps* function (packages GenomicAlignments of R) to generate the matrix of read count for each gene from BAM files. Differentially expressed genes (DEG) lists for *CsDML* OX vs. WT were obtained using the function exactTest (edgeR package of R) (Robinson et al., 2010). For DEG analyses, *CsDML* OX1 and *CsDML* OX3 transcriptomes were considered as biological replicates when comparing to WT, using a *FDR* < 0.1. Gene annotations for DEG were obtained from the *Populus trichocarpa* annotation table v.3.0. http://www.phytozome.net. Gene ontology (GO) categories were analyzed using the web-based platform agriGO at http://bioinfo.cau.edu.cn/agriGO (Du et al., 2010). Enriched GO categories were identified using an FDR-adjusted value of ≤ 0.05 as the cutoff for significance.

## Author contributions

D.C., P.G-M. and I.A. planned and designed the research; D.C, A.M-C., JM. R-S. and C. D. performed experiments; D.C., M.P., P. G-M. and I.A. analyzed data; D.C., M.K., M.P., P.G-M. and I.A. wrote the manuscript.

## Supporting information

The following materials are available in the online version of this article.

**Figure S1.**
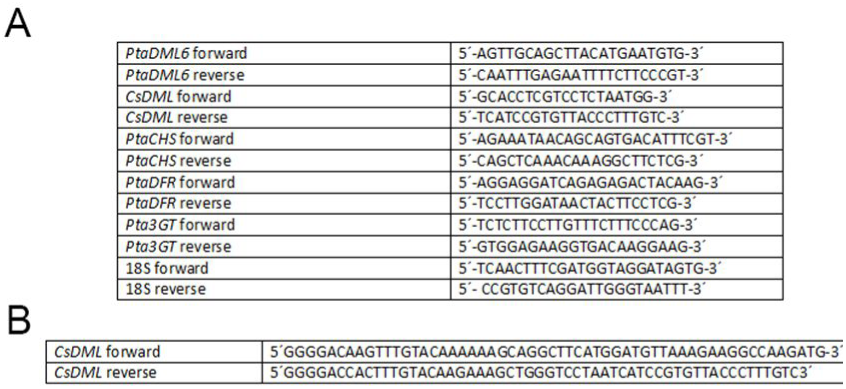
List of primers used in this work. List of primers used in the present study. *A*, List of primers used for the analyses of gene expression by RT-PCR. B, Primers used for amplification the CDS of *CsDML* gene with attB sites to generate the constructions needed for the overexpression of *CsDML* in poplar.

**Figure S2.**
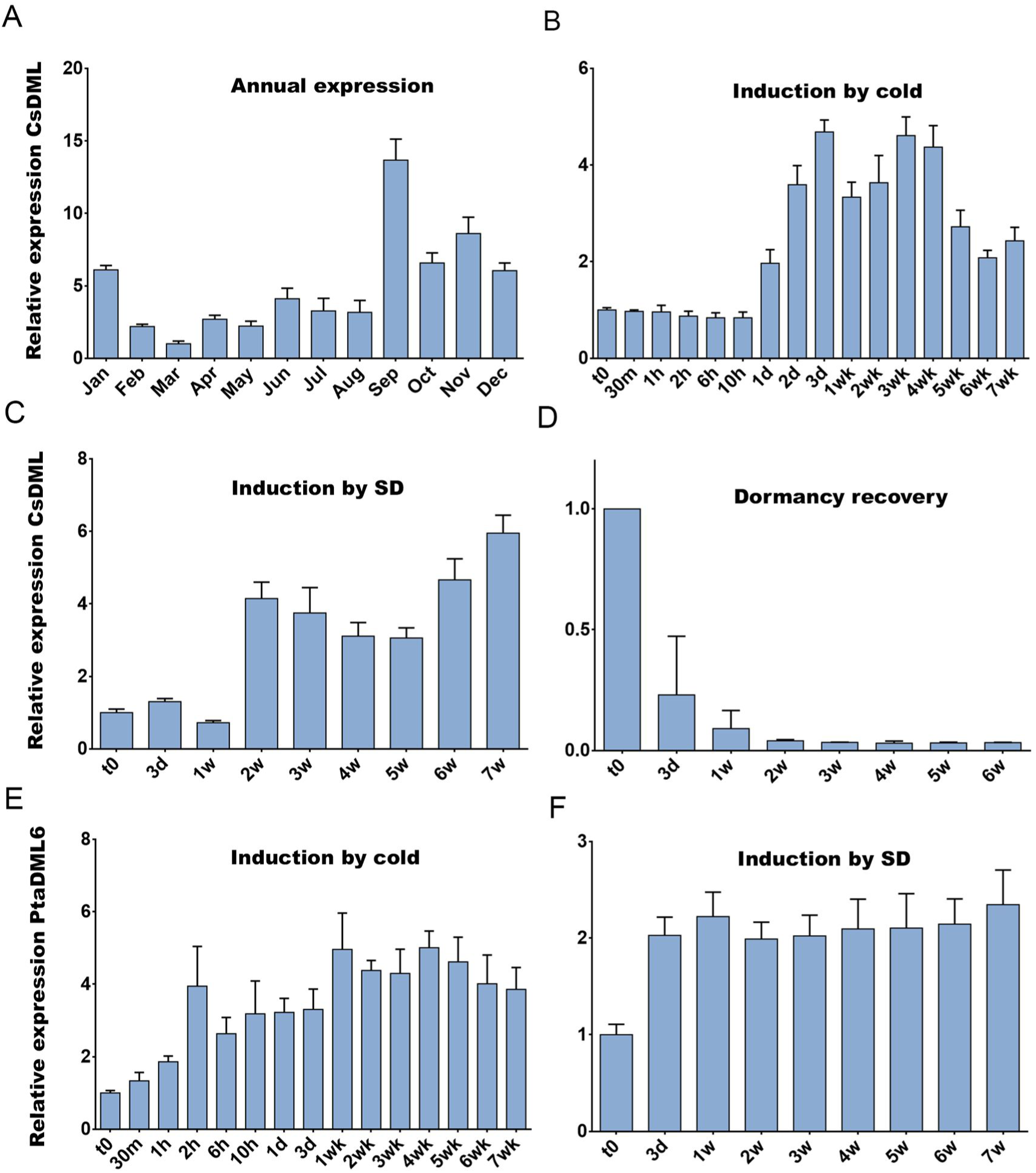
Characterization of CsDML gene expression in chestnut trees. Quantitative RT-PCR analysis of: *A, CsDML* annual expression pattern in 2-year-old chestnut branches from adult trees growing under natural conditions; B, *CsDML* cold induction expression in chestnut stems from 8-months-old plants grown in a controlled-environment growth chamber (LD, 22°C) and subsequently transferred to LD, 4°C for 7 weeks; C, *CsDML* SD induction expression in chestnut stems from 8-month-old plants grown in a controlled-environment growth chamber (LD, 22°C) and subsequently transferred to SD, 22°C for 6 weeks; D, *CsDML* expression levels in chestnut stems from 11-month-old endodormant plants grown under natural conditions, and transferred before chilling fulfillment to a controlled-environment growth chamber in standard conditions (LD, 22 °C); E, *PtaDML6* cold induction expression in poplar stems from 8-weeks-old plants grown in a controlled-environment growth chamber (LD, 22°C) and subsequently transferred to LD, 4°C for 7 weeks; F, *PtaDML6* SD induction expression in poplar stems from 6-weeks-old plants grown in a controlled-environment growth chamber (LD, 22°C) and subsequently transferred to SD, 22°C for 7 weeks. In A,B,C,D,E,F plotted values and error bars are the fold-change means ± SD recorded in three technical replicates. In D, plotted values and errors bars are the fold-change means ± SD recorded in two biological replicates.

**Figure S3.**
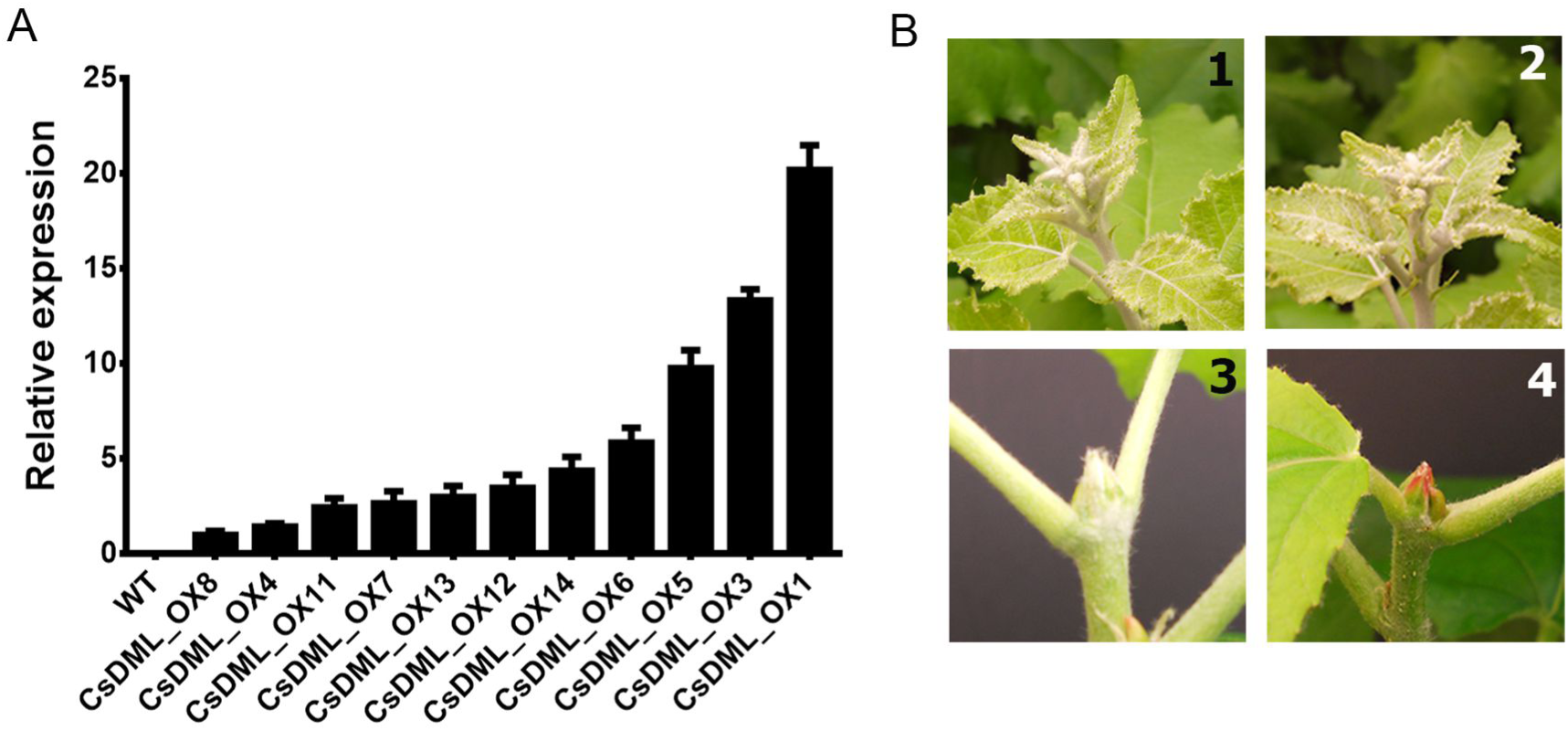
Characterization of the CsDML OX overexpressing poplar trees. A, quantitative RT-PCR analysis of *CsDML* in the *CsDML* OX (line OX1) transgenic plantlet leaves. Samples were obtained from plantlets grown in a greenhouse for 5 weeks in conditions of LD, 22°C. Plotted values and error bars are the fold-change means ± SD recorded in three technical replicates. B, B1 and B2 indicate WT and *CsDML* OX (line OX1) plants after growth for 4 weeks in conditions of LD,22°C. At this point no phenotypical differences, compared to WT, were observed. B3 and B4 indicate WT and *CsDML* OX (line OX1) plants after growth for 49 days in conditions of SDs, 19°C. At this point, bud maturation was advanced in *CsDML* OX compared to the WT. The same occurs for the other OX line.

**Figure S4.**
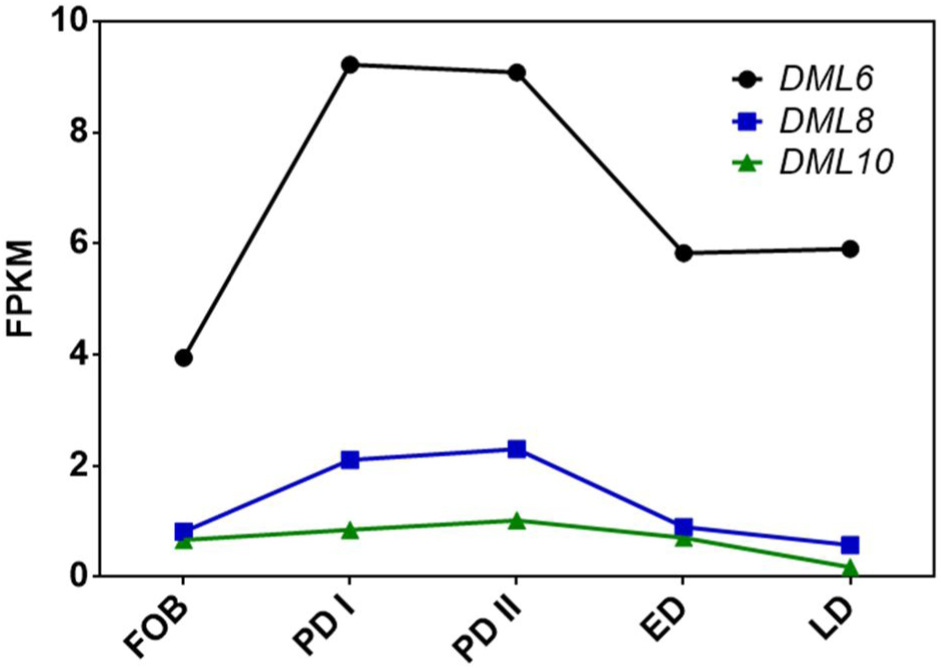
Expression of poplar DMLs in apex. FPKM values of *PtDML6, PtDML8* and *PtDML10* in public RNA seq analyses performed in *Populus trichocarpa* apex during different developmental stages along the year; FOB: fully open bud; PDI-PDII: predormant bud I and II; ED: early dormant bud; LD: late dormant bud. These data are available in phytozome11 website (https://phytozome.jgi.doe.gov/pz/portal.html).

**Table S1.** List of DEG in *CsDML* OX in comparison to WT.

**Table S2.** Ontology classification of downregulated genes in CsDML OX apices.

## ACKNOWLEDGMENTS

The authors thank C. Aragoncillo, J. Matus, and J. Paz-Ares for helpful suggestions on the manuscript. The authors also thank R. Iglesias and M.V. Pérez Chaca for technical advice and assistance with the fluorescence *in situ* hybridization experiments.

